# TManual: Assistant for manually measuring length development in structures built by animals

**DOI:** 10.1101/2022.12.21.521503

**Authors:** Nobuaki Mizumoto

## Abstract

1. Structures built by animals are extended phenotypes, and animal behavior can be better understood by recording the temporal development of structure construction. For most subterranean and wood-boring animals, these structures consist of gallery systems, such as burrows made by mice, tunnel foraging by termites, and nest excavation in ants. Measurement of the length development of such structures is often performed manually. However, it is time-consuming and cognitively costly to track length development in nested branching structures, hindering the quantitative determination of temporal development.
2. Here, I introduce TManual, which aids the manual measurement of structure length development using a number of images. TManual provides a user interface to draw gallery structures and take over all other processes handling input datasets (e.g., zero-adjustment, scaling the units, measuring the length, assigning gallery identities, and extracting network structures). Thus, users can focus on the measuring process without interruptions.
3. As examples, I provide the results of the analysis of a dataset of tunnel construction by three termite species after successfully processing 1,125 images in ∼3 h. The output datasets clearly visualized the interspecific variation in tunneling speed and branching structures. Furthermore, I applied TManual to a complex gallery system by another termite species and extracted network structures.
4. Measuring the lengths of objects from images is an essential task in biological observation. TManual helps users handle many images in a realistic time scale, enabling a comparative analysis across a wide array of species. TManual does not require programming skills and outputs a tidy data frame in CSV format. Therefore, it is suitable for any user who wants to perform image analysis for length measurements.

## Introduction

Structures built by animals are considered extended phenotypes, and the temporal development of such structures reflects the temporal dynamics of the animal’s behavior (Hansell, 2005; Sugasawa and Pritchard, 2022). One common type of structure built by animals is a gallery system, which is observed in most subterranean animals, and includes burrow construction in mice (Bedford *et al*., 2022; Metz *et al*., 2017), tunnel foraging by termites (Bardunias and Su, 2005; Mizumoto *et al*., 2020), and nest excavation in ants (Buhl *et al*., 2005; Toffin *et al*., 2009). Gallery systems can also be observed aboveground in some social insects, for example, shelter-tube construction by ants and termites (Chiu *et al*., 2022; Mizumoto and Bourguignon, 2020). Tracking the temporal development of gallery structures is important for understanding the dynamics of collective nest building.

Temporal development is often observed in two-dimensional experimental setups (e.g., Figures 1-3), and researchers have used two different approaches to capture the geometric properties of structures. Some studies have focused on overall structural patterns, such as the excavated area (Buhl *et al*., 2005), perimeter (Mizumoto *et al*., 2015; Toffin *et al*., 2010, 2009), and nodes in a gallery system (Buhl *et al*., 2004; Gravish *et al*., 2012; Perna *et al*., 2008). The main advantage of this approach is that it can be automated, with the parameters automatically obtained by image-processing programs after binarizing them (which is often achieved using program languages such as C++ software). However, this approach requires high-quality standardized recording setups, and the interpretation of outcomes is sometimes not intuitive. For example, several different morphological patterns can increase/decrease the perimeter of a gallery system.

**Figure 1.**
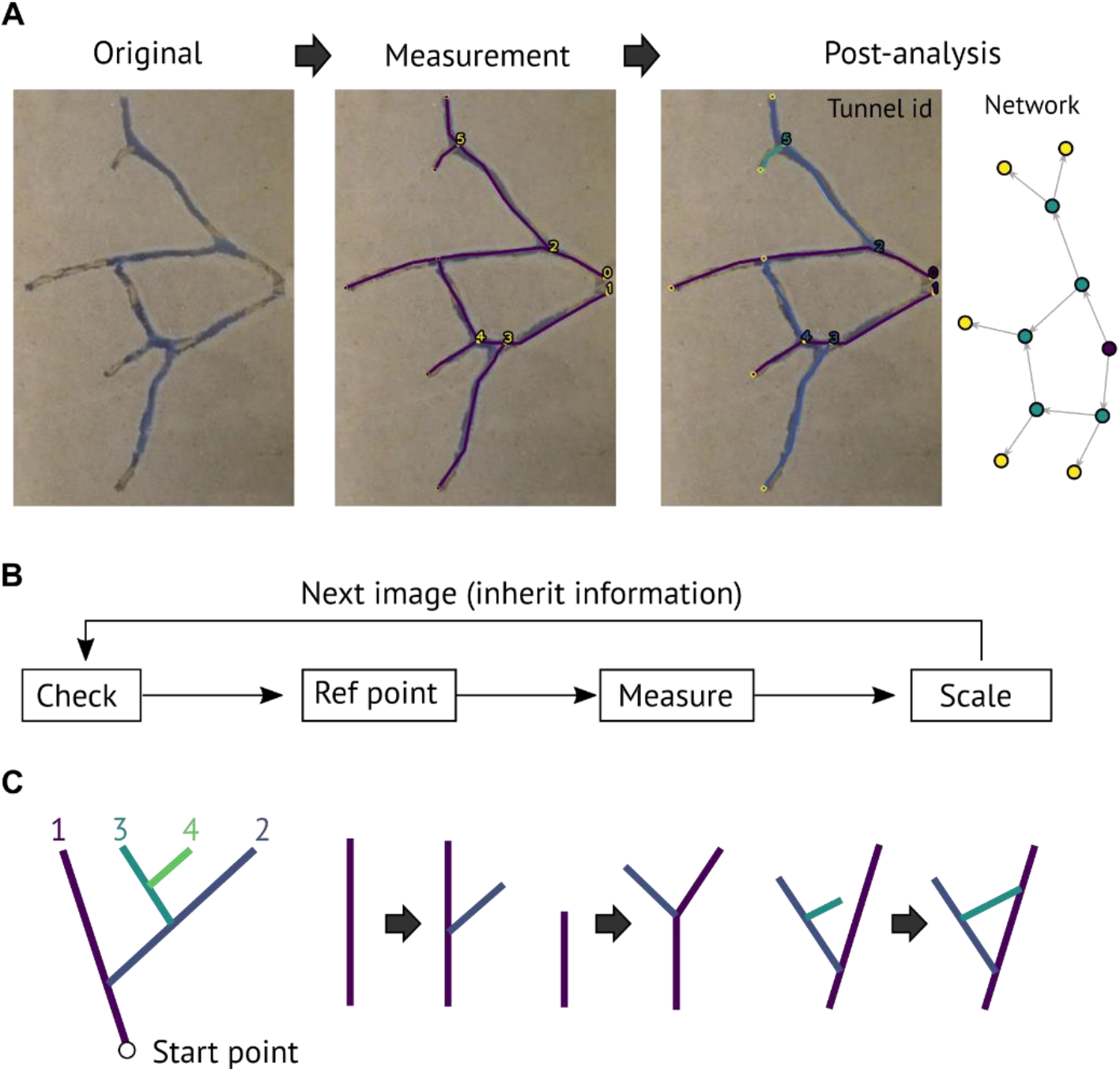
Overview of the TManual. TManual aids manual measurement of the temporal development of gallery-formed structures. (A) An overview of the image-processing procedure. First, the user draws all gallery structures by clicking on the image. Then the post-analysis program automatically categorizes galleries (primary, secondary, tertiary, …, in different colors), calculates gallery length with scaling in mm, reconstruct network structures, and outputs tidy data frames in CSV format for subsequent analysis. (B) Overflow of the measurement process. TManual sequentially shows images and asks users to analyze or skip the image (Check), identify a consistent landmark as a reference if required (Ref point), draw galleries in a freeform object with straight segments (Measure), and select scale objects (Scale). The information will be inherited in the next image. (C) The rule used to identify tunnel identity.

Most previous studies have directly measured the length of each gallery in a whole system and categorized them into different groups to capture the geometric patterns. This approach has been adopted to assess ant nest excavation (Kwapich *et al*., 2018; O’Fallon *et al*., 2022), termite tunnel foraging (Hedlund and Henderson, 1999; Mizumoto *et al*., 2020; Robson *et al*., 1995; Su *et al*., 2004), shelter-tube construction (Mizumoto and Matsuura, 2013), and cooperative burrowing in mice (Bedford *et al*., 2022). In this approach, because experimenters arbitrarily identify and measure each gallery, the outcomes are intuitive and the results are easy to interpret. However, because this process requires considerable effort, most studies have only focused on one or a few snapshots of structural development. This hinders our understanding of the dynamics of gallery-building behavior and prevents a comparative analysis across taxa. Still, a manual analysis can be useful for handling images with much noise (e.g., it is common for images of structures to contain non-structure objects, such as excavated substrates and individuals). Thus, it is important to develop a simpler way to aid the manual measuring process and to effectively process a large number of images.

The key stages in the manual process include measuring gallery length, identifying nodes, assigning galleries into categories, measuring the length of an object for scaling, and storing the information obtained in an organized file format for subsequent data analysis. This often requires users to move back-and-forth between image analysis software and spreadsheets, which makes the process labor-intensive and cognitively costly; it can also result in unintentional human errors. To overcome these problems, I herein introduce the TManual software, which is designed to achieve stress-free and quick manual measurement of gallery length. The user simply needs to click on the points of interest in images to obtain spreadsheets that include all geometric information.

## Software

TManual is based on Python3, and the source code is released under the BSD-3 License at https://github.com/nobuaki-mzmt/tmanual. The EXE file is also available for Microsoft Windows users. The future development of the software will also be announced in this GitHub deposit. TManual consists of two processes: measurement and post-analysis. The user should prepare the sequential image files (e.g., JPG), which should be named “id_serial,” e.g., [TunnelA_00.jpg, TunnelA_01.jpg, TunnelA_02.jpg, …, TunnelA_19.jpg, TunnelA_20.jpg, TunnelB_00.jpg, TunnelB_01.jpg, …].

## Measurement

The measurement program is designed to display all images of interest sequentially and store all user inputs. It works according to the process outlined below (Fig. 1):

### 1. Check

TManual displays the image together with an overlay of the gallery information of the previous image (or the current image if it is already analyzed). If users do not find any development in the gallery system compared to the previous image they can skip the image, and TManual copies the information of the previous image to the current one. Similarly, if users find no gallery in the image, TManual generates data with 0 values for subsequent analysis.

### 2. Ref point

TManual asks the user to indicate the reference point, which is an identifiable landmark across all images (e.g., the corner of the experimental arena). This is useful when the relative position of the camera and object is not fixed (e.g., when users take photos every 24 h and need to bring the experimental arena under the camera when filming). If the camera and object are fixed, users can skip this process (the reference point will be the top left corner of the image).

### 3. Measure

Users draw the galleries as freeform line objects with straight segments. For branching structures, two gallery lines need to contact (< threshold pixel, users can decide). The start and end points of galleries are treated as nodes to reconstruct a network structure (nodes within the threshold are regarded as the same node). If the users follow the gallery identity definition in the next section (also in Fig. 1C), each gallery is assigned to one of the categories (primary, secondary, tertiary, …).

### 4. Scale

Measure the length of the scale object. This is used to convert the unit from pixels to mm during the post-analysis stage.

All the user inputs are stored in res.pickle, and are used in the following post-analysis stage.

### Post-analysis

The post-analysis program creates CVS files containing all of the information about the gallery structures based on res.pickle. This includes the length of each gallery (and total length), the number of galleries, the number of nodes, gallery classification, and network structure of gallery system.

In termite foraging tunnels, many studies have categorized branching tunnels into primary, secondary, tertiary, and quaternary tunnels to capture the geometry of gallery structures (Hedlund and Henderson, 1999; Su *et al*., 2004). All tunnels originating from the initial point are classified as primary, and primary tunnels are often extended to the farthest point possible (Hedlund and Henderson, 1999). Tunnels branching from the primary tunnel are classified as secondary tunnels; tertiary tunnels branch from secondary tunnels; and quaternary tunnels branch from tertiary tunnels. However, this definition cannot always determine tunnel identity uniquely (Su *et al*., 2004); e.g., some researchers only focus on one snapshot, and tunnel identity can change according to temporal developments (Hedlund and Henderson, 1999). Hence, expanding the definition of previous studies, I defined tunnels as follows:

1. Primary tunnels originate from the start point.
2. Tunnels that emerge from the side of preexisting tunnels are descendant tunnels (primary -> secondary, secondary -> tertiary, tertiary -> quaternary, and following).
3. If one tunnel splits into two branches, the one with a shallower angle retains the original identity, while the other is a descendant tunnel.
4. The point where a tunnel merges with a preexisting tunnel is considered the endpoint.

Users should follow this definition during the above *Measurement* process. Then the post-analysis program automatically assigns identities for each gallery and measures the total length and number of each category. By indicating the length of the scale object measured in the *Measurement* program, all outcomes are scaled in the unit of mm. The *post-analysis* program outputs images overlaying user-labeled gallery structures with gallery identities. These images can be used to confirm if the post-analysis works as users intended and to correct specific images if required.

### Examples of application

#### Comparative analysis of tunnel time development in termites

To demonstrate the application of the software, I analyzed the development of termite foraging tunnels observed in a previous study (Mizumoto *et al*., 2020). The previous study measured tunnel development by three different termite species (*Paraneotermes simplicicornis, Reticulitermes tibialis*, and *Heterotermes aureus*) in a two-dimensional experimental arena. Although the recording was performed for 24 h, this study only analyzed one snapshot for each of the 46 experiments. This is because that study used the generic software ImageJ (National Institutes of Health, Bethesda, MD) and avoided the time-consuming and labor-intensive manual process of measuring tunnel length and structures for all snapshots. I overcame this problem using TManual. I analyzed the tunnel structures every hour in 45 experiments, resulting in an analysis of 25 × 45 = 1,125 images.

It took ∼3 h to analyze all of the images using TManual (all analyzed images are available at Video S1). Even with the manual process, the processing speed of ∼400 images/h was sufficiently cost-effective to produce high-resolution datasets for assessing temporal development. However, it should be noted that the required time will be based on the complexity of the tunnel structures and size, and the analysis will take longer for gallery structures with many nested branching patterns.

A previous study found interspecific variation in excavation speed and tunnel structures from snapshots (Mizumoto *et al*., 2020). Applying TManual, I successfully reproduced and better quantified the results, particularly the visualization of temporal development. First, it was found that *H. aureus* and *R. tibialis* excavated longer tunnels than *P. simplicicornis* (Fig. 2A), consistent with the previous observation that the former two species reached the arena wall faster than the latter (Mizumoto *et al*., 2020). The tunnel identification provided by TManual captured the interspecific variation in tunnel geometry (Fig. 2BC). It was previously indicated that *H. aureus* builds more branching tunnels than *R. tibialis* and *P. simplicicornis*, which was confirmed by counting the number of ends of tunnels (Mizumoto *et al*., 2020). TManual quantified this variation by showing that *H. aureus* builds more secondary tunnels than *R. tibialis* and *P. simplicicornis*. Despite *R. tibialis* excavating longer tunnels, *P. simplicicornis* and *R. tibialis* produced very similar structures.

**Figure 2.**
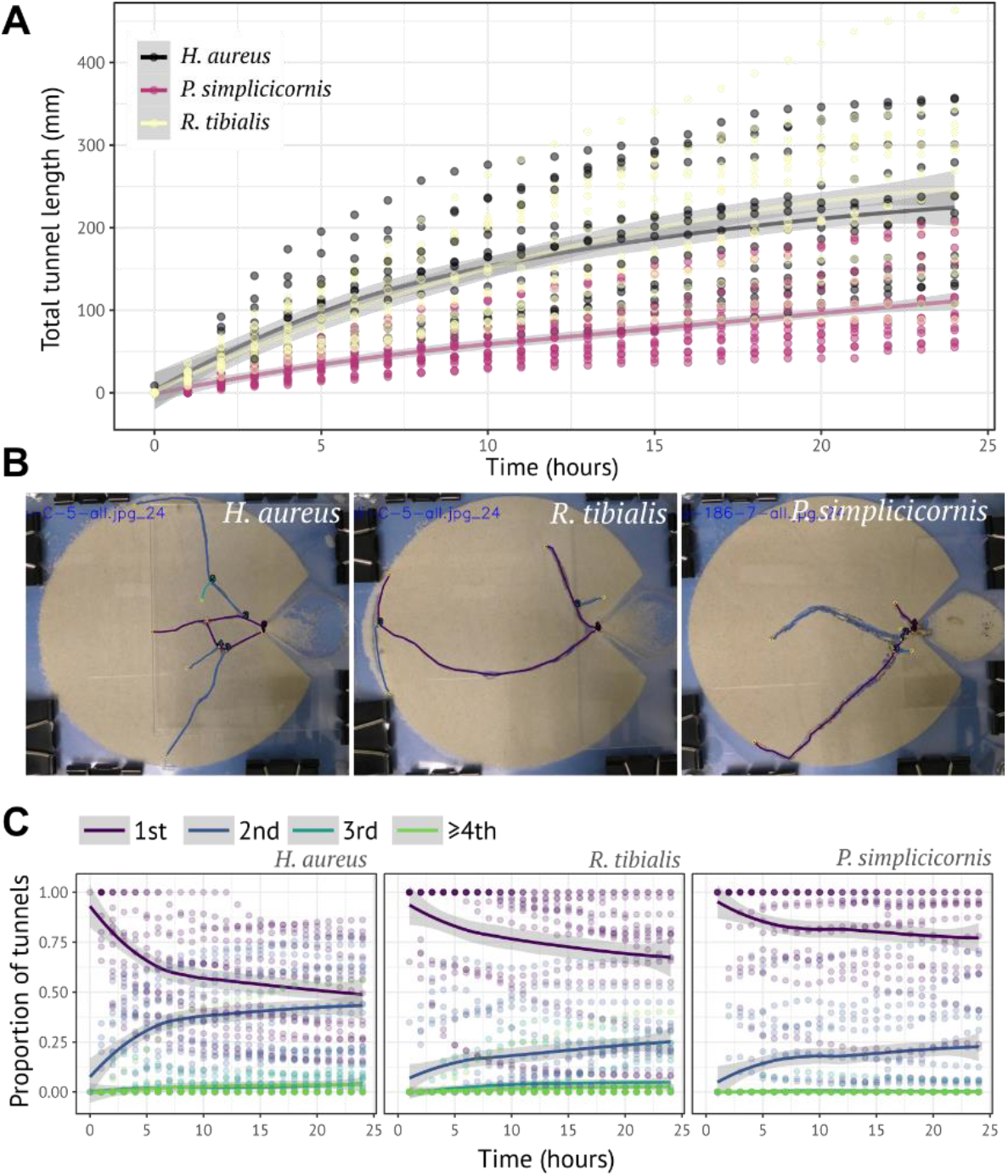
An example of the application of TManual. The results of analysis of 1,125 images of termite tunnel development by TManual. (A) The temporal development of total tunnel length. *Heterotermes aureus* and *Reticulitermes tibialis* built longer tunnels than *Paraneotermes simplicicornis*. (B) Representative images analyzed by TManual. The gallery structures with tunnel identity were overlayed on the original images provided by the *post-analysis* program. (C) Comparison of tunnel geometry between species. 1st, 2nd, 3rd, and ≥4th indicate primary, secondary, tertiary, and quaternary or later tunnels. The structures are described by the proportion of each tunnel identity, with *H. aureus* building more branching tunnels than the other species.

These results demonstrate that TManual is a cost-effective and strong tool for the manual measurement of gallery development by animals.

#### Network structure of termite foraging tunnels

In the above example, gallery structures were described by tunnel identities. However, for more complex structures, another way is to regard the gallery system as edges and nodes that form a network structure. The *post-analysis* program of TManual reconstructs network structures from starting and ending points of each gallery drawn by users. As examples, I analyzed two foraging tunnel structures built by a termite, *Coptotermes formosanus* (Fig. 3). TManual successfully reconstructed the foraging tunnel networks in termites, one with 374 nodes and 535 edges (Fig. 3A), and the other with 116 nodes and 140 edges (Fig. 3B).

**Figure 3.**
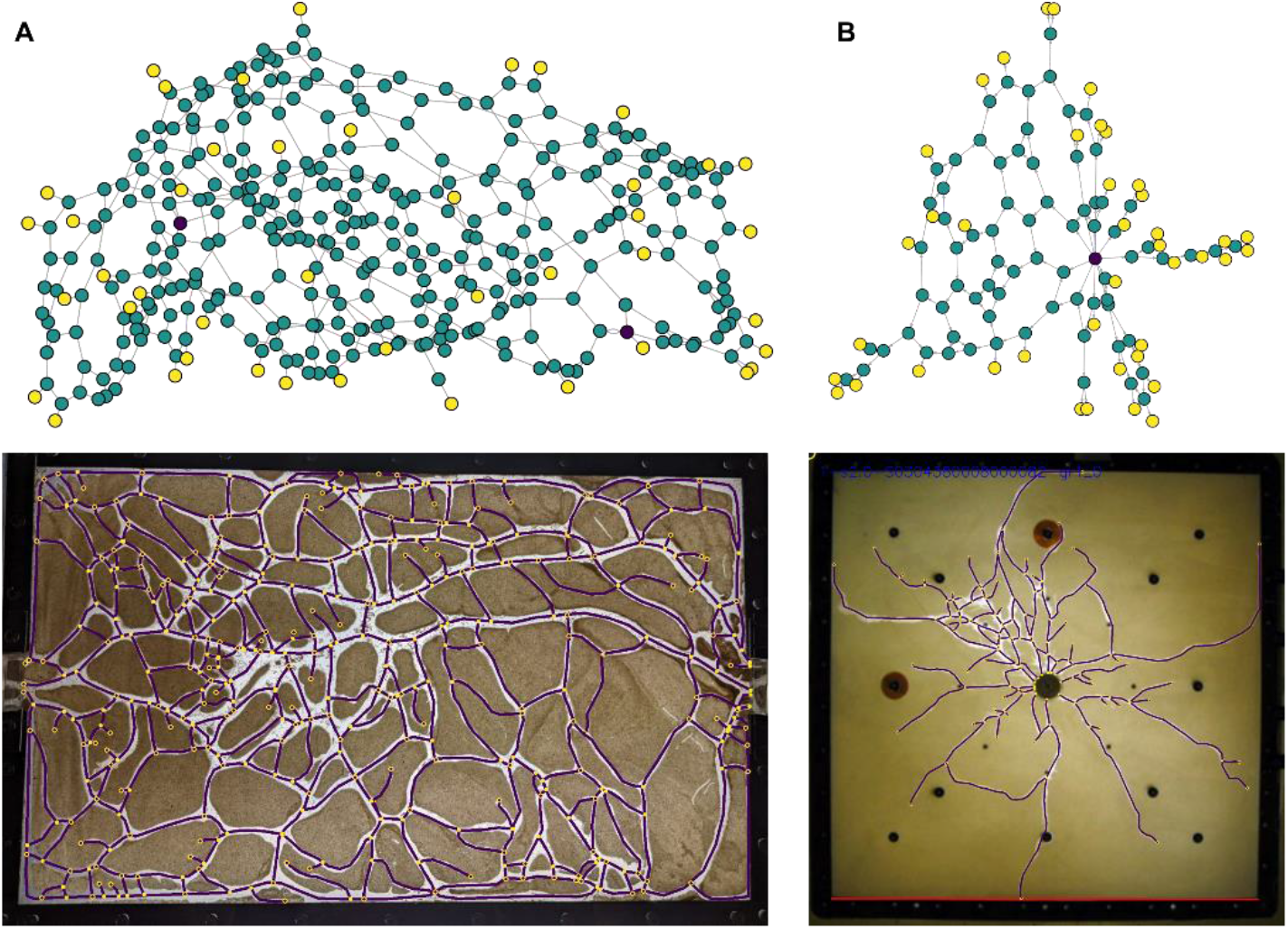
Network structures provided by post-analysis program in TManual. Tunnel network structures created by *C. formosanus* termites in two-dimensional arenas. Purple nodes indicate entrance, green nodes indicate intersection, and yellow nodes indicate tunnel ends. (A) The structure created in arena between two entrances. Photo is provided by Sang-Bin Lee. (B) Foraging tunnel structures created from a single entrance in the center. The photo used in (Lee et al., 2008) was used for the analysis.

## Future developments

TManual assumes that galleries become longer according to time and does not support the case when galleries become shorter. For example, during the tunneling process, termites often backfill already-excavated tunnels. Treating this backfilling process might depend on the user’s interpretation. For example, if users are interested in the excavated distances during the observations rather than the final structures, they need not be concerned about the backfilling process. If users are interested in the outcome structures at each time point, a reanalysis is required when backfilling occurs. TManual can handle this by selecting the “not appending” option in the *Check* process of *Measurement*. However, the easiness of this process could be improved with further updates.

## Conclusion

Measuring the lengths of objects from images is a basic task in biological observations and this task needs to be completed in a quick and simple way. TManual provides a user interface to manually measure the length development from multiple images. Although I present the software with a focus on the termite tunneling system, it can be applied to similar structures built by other animals, such as ant nest excavations and burrows dug by mice. Furthermore, as shown by studies of root development in plants (Kume *et al*., 2018), branching structures are ubiquitous in nature, which provides another potential application of TManual.

## Acknowledgments

I thank Sang-Bin Lee for informing me that this tool will likely be helpful for other researchers, encouraging me to write this paper, and providing several sample images; Kaitlin Gazdick for assistance in designing the software specifications, performing several test runs, and providing several sample images; and Jamie M. Kass for advice in depositing source codes. This study was supported by a JSPS Research Fellowships for Young Scientists CPD (grant number: 20J00660).

## Data accessibility statement

I will archive the source codes of TManual in Zenodo upon acceptance. Updated source codes can be accessed at the GitHub repository: https://github.com/nobuaki-mzmt/tmanual/.

## References

Bardunias P, Su NY. 2005. Comparison of tunnel geometry of subterranean termites (Isoptera: Rhinotermitidae) in “two-dimensional” and “three-dimensional” arenas. Sociobiology 45:679–685.

Bedford NL, Weber JN, Tong W, Baier F, Kam A, Greenberg RA, Hoekstra HE. 2022. Interspecific variation in cooperative burrowing behavior by Peromyscus mice. Evolution Letters 6:330–340. doi:10.1002/evl3.293

Buhl J, Deneubourg J-L, Grimal A, Theraulaz G. 2005. Self-organized digging activity in ant colonies. Behavioral Ecology and Sociobiology 58:9–17. doi:10.1007/s00265-004-0906-2

Buhl J, Gautrais J, Deneubourg JL, Theraulaz G. 2004. Nest excavation in ants: Group size effects on the size and structure of tunneling networks. Naturwissenschaften 91:602–606. doi:10.1007/s00114-004-0577-x

Chiu C, Chen B, Chang F, Kuan K, Hou-Feng L. 2022. Functional plasticity of foraging shelter tubes built by termites. Environmental Entomology 51:649–659. doi:10.1093/ee/nvac054

Gravish N, Garcia M, Mazouchova N, Levy L, Umbanhowar PB, Goodisman MAD, Goldman DI. 2012. Effects of worker size on the dynamics of fire ant tunnel construction. Journal of The Royal Society Interface 9:3312– 3322. doi:10.1098/rsif.2012.0423

Hansell MH. 2005. Animal architecture. Oxford: Oxford University Press.

Hedlund JC, Henderson G. 1999. Effect of available food size on search tunnel formation by the Formosan subterranean termite (Isoptera: Rhinotermitidae). Journal of Economic Entomology 92:610–616. doi:10.1093/jee/92.3.610

Kume T, Ohashi M, Makita N, Kho LK, Katayama A, Endo I, Matsumoto K, Ikeno H. 2018. Image analysis procedure for the optical scanning of fine-root dynamics: Errors depending on the observer and root-viewing window size. Tree Physiology 38:1927–1938. doi:10.1093/treephys/tpy124

Kwapich CL, Valentini G, Hölldobler B. 2018. The non-additive effects of body size on nest architecture in a polymorphic ant. Philosophical Transactions of the Royal Society B: Biological Sciences 373:20170235. doi:10.1098/rstb.2017.0235

Lee S-H, Bardunias PM, Su N-Y. 2008. Two strategies for optimizing the food encounter rate of termite tunnels simulated by a lattice model. Ecological Modelling 213:381–388. doi:10.1016/j.ecolmodel.2008.01.004

Metz HC, Bedford NL, Pan YL, Hoekstra HE. 2017. Evolution and genetics of precocious burrowing behavior in Peromyscus mice. Current Biology 27:3837-3845.e3. doi:10.1016/j.cub.2017.10.061

Mizumoto N, Bardunias PM, Pratt SC. 2020. Complex relationship between tunneling patterns and individual behaviors in termites. American Naturalist 196:555–565. doi:10.1086/711020

Mizumoto N, Bourguignon T. 2020. Modern termites inherited the potential of collective construction from their common ancestor. Ecology and Evolution 10:6775–6784. doi:10.1002/ece3.6381

Mizumoto N, Kobayashi K, Matsuura K. 2015. Emergence of intercolonial variation in termite shelter tube patterns and prediction of its underlying mechanism. Royal Society Open Science 2:150360. doi:10.1098/rsos.150360

Mizumoto N, Matsuura K. 2013. Colony-specific architecture of shelter tubes by termites. Insectes Sociaux 60:525–530. doi:10.1007/s00040-013-0319-1

O’Fallon S, Lowell ESH, Daniels D, Pinter-Wollman N. 2022. Harvester ant nest architecture is more strongly affected by intrinsic than extrinsic factors. Behavioral Ecology 33:644–653. doi:10.1093/beheco/arac026

Perna A, Jost C, Couturier E, Valverde S, Douady S, Theraulaz G. 2008. The structure of gallery networks in the nests of termite Cubitermes spp. revealed by X-ray tomography. Naturwissenschaften 95:877–884. doi:10.1007/s00114-008-0388-6

Robson SK, Lesniak MG, Kothandapani RV, Traniello JFA, Thorne BL, Fourcassié V. 1995. Nonrandom search geometry in subterranean termites. Naturwissenschaften 82:526–528. doi:10.1007/BF01134490

Su NY, Stith BM, Puche H, Bardunias P. 2004. Characterization of tunneling geometry of subterranean termites (Isoptera: Rhinotermitidae) by computer simulation. Sociobiology 44:471–483.

Sugasawa S, Pritchard DJ. 2022. The significance of building behavior in the evolution of animal architecture. Ecological Research 1–9. doi:10.1111/1440-1703.12309

Toffin E, Di Paolo D, Campo A, Detrain C, Deneubourg JL. 2009. Shape transition during nest digging in ants. Proceedings of the National Academy of Sciences of the United States of America 106:18616–18620. doi:10.1073/pnas.0902685106

Toffin E, Kindekens J, Deneubourg J-L. 2010. Excavated substrate modulates growth instability during nest building in ants. Proceedings of the Royal Society of London B 277:2617–25. doi:10.1098/rspb.2010.0176

